# Dissecting phylogenetic signal and accounting for bias in whole-genome data sets: a case study of the Metazoa

**DOI:** 10.1101/013946

**Authors:** Marek L. Borowiec, Ernest K. Lee, Joanna C. Chiu, David C. Plachetzki

## Abstract

Transcriptome-enabled phylogenetic analyses have dramatically improved our understanding of metazoan phylogeny in recent years, although several important questions remain. The branching order near the base of the tree is one such outstanding issue. To address this question we assemble a novel data set comprised of 1,080 orthologous loci derived from 36 publicly available genomes and dissect the phylogenetic signal present in each individual partition. The size of this data set allows for a closer look at the potential biases and sources of non-phylogenetic signal. We assessed a range of measures for each data partition including information content, saturation, rate of evolution, long-branch score, and taxon occupancy and explored how each of these characteristics impacts phylogeny estimation. We then used these data to prepare a reduced set of partitions that fit an optimal set of criteria and are amenable to the most appropriate and computationally intensive analyses using site-heterogeneous models of sequence evolution. We also employed several strategies to examine the potential for long-branch attraction to bias our inferences. All of our analyses support Ctenophora as the sister lineage to other Metazoa, although support for this relationship varies among analyses. We find no support for the traditional view uniting the ctenophores and Cnidaria (jellies, anemones, corals, and kin). We also examine phylogenetic placement of myriapods (centipedes and millipedes) and find it more sensitive to the type of analysis and data used. Our study provides a workflow for minimizing systematic bias in whole genome-based phylogenetic analyses.

## Introduction

Advances in sequencing technology have led to a revolution in genomics, where draft genome assemblies for most species can be obtained at relatively low cost. One of the most significant outcomes anticipated of this revolution is an understanding of the interrelationships of the major lineages of multicellular animals, the Metazoa. A robust phylogeny for Metazoa will provide evolutionary context allowing a narrative on the timing and origins of the major features of animals including nervous systems (Moroz et al. 2014), immune systems (Bosch et al. 2009), cell types (Arendt 2008) and other complex traits. This phylogenetic framework will also provide important insights into the role that convergence could play in the evolution of these traits. Here we approach the question of metazoan relationships by dissecting the phylogenetic signal present in a novel dataset derived from 36 publicly available whole genome sequences. We focus our analyses on identifying and ameliorating potential sources of bias that could stem from the inclusion of long-branch taxa or from data partitions with specific bias-inducing properties. We also explore conflicting signal between alternative modes of phylogenetic analysis.

To date, numerous studies have applied large sequence datasets, drawn mostly from transcriptome sequencing efforts, to the problem of metazoan phylogeny (Dunn et al. 2008; Hejnol et al. 2009; Philippe et al. 2009; Pick et al. 2010; Nosenko et al. 2013). Such approaches have yielded several important findings in recent years. First, the phylogenetic position of the comb jellies (phylum Ctenophora) as sister to all remaining metazoan phyla including sponges (phylum Porifera) has been proposed in several recent analyses. This surprising finding has attracted much attention because it suggests that neurons and other complex traits, present in ctenophores and eumetazoans but absent in sponges or placozoans, either evolved twice in Metazoa or were independently, secondarily lost in the lineages leading to sponges and placozoans (Ryan et al. 2013; Moroz et al. 2014; Ryan 2014; Dunn et al. 2014). This relationship was first suggested by phylogenetic analyses of transcriptome datasets (Dunn et al. 2008) and later by similar analyses that were augmented by whole genome sequences of two additional ctenophore species; *Mnemiopsis leidyi* (Ryan et al. 2013) and *Pleurobrachia bachei* (Moroz et al. 2014). However, this finding is controversial and several alternative analyses have suggested that the basal position of ctenophores could be the result of long-branch attraction (LBA) or other artifacts stemming from noise present in large alignments (Philippe et al. 2009; Pick et al. 2010; Philippe et al. 2011). In addition, the position of the centipedes and millipedes (Myriapoda) relative to other arthropods has also come into focus recently through transcriptome-based analyses (Rota-Stabelli et al. 2011; Rehm et al. 2013). These studies argue for the placement of myriapods as sister to insects and crustaceans (Pancrustacea), a finding that is consistent with several morphological features shared between the two groups (Rota-Stabelli et al. 2011). However, other hypotheses for the placement of myriapods are common in phylogenomic studies (Dunn et al. 2008; Hejnol et al. 2009; Meusemann et al. 2010).

While transcriptome-enabled phylogenetic analyses have doubtlessly proven powerful in the fabrication of large datasets representing large numbers of taxa, several caveats to this approach deserve mention. First, phylogenetic datasets derived in this way can only capture information on genes that are expressed in the tissue collected for a given taxon. Whole organism transcriptome datasets are possible for small organisms, but many taxa can only be represented by transcriptomes from selected tissues. The incomplete nature of transcriptome data is compounded when considering taxa with complicated life histories, which account for the majority of metazoan taxa. In addition, most transcriptome data derived from short read, next generation sequencing approaches, require assembly prior to their use in phylogenetics. Since many popular assembly procedures take into account the quantitative prevalence of specific transcripts in raw short read data (Hass et al. 2013), the transcriptome models that are produced by these procedures are a function of gene expression level, which can differ between taxa and tissues. Further, genes that are lowly expressed may not be recovered in transcriptome assembly. Each of these factors could impose biases on the types of genes procured by these procedures for phylogenetic analyses. In addition, since orthologous gene prediction for phylogenetic analyses aims to maximize data coverage, both in terms of taxon occupancy and in the number of loci, biases in transcriptome assembly in a few taxa can lead to the exclusion of informative partitions from the analysis. Matrices constructed in this way are also often sparse, consisting of much missing data, which can confound phylogenetic analyses (Lemmon et al. 2009; Roure et al. 2013). Finally, transcriptome datasets have been occasionally shown to include contaminants from other taxa, which could disrupt accurate phylogenetic reconstruction (Philippe et al. 2011). Whole genome sequences, while not without drawbacks of their own, do offer a solution to many of the above issues encountered in transcriptome-based phylogenetic analyses.

The purpose of this study is to examine metazoan phylogeny with a focus on recent controversies surrounding the rooting of the animal tree and the positions of the ctenophores and myriapods using an alternative data set obtained exclusively from taxa with publicly available whole genome sequences. While previous studies of metazoan phylogeny included matrices derived from whole genome sequences (Srivastava et al. 2010; Ryan et al. 2013), the data set compiled here is by far the largest in terms of number of characters and taxa. Our novel data set is drawn from the gene models of 34 metazoan and two choanoflagellate genomes (supplementary table 1). We use a highly accurate orthology prediction procedure followed by stringent alignment filtering to recover 1,080 phylogenetically informative orthologous groups (OGs) that bear on metazoan phylogeny. We assessed a range of measures for each data partition including information content, saturation, rate of evolution, long-branch score, and taxon occupancy and explored how each of these characteristics impacts phylogeny estimation. We then used these data to prepare a reduced set of partitions that fit an optimal set of criteria and are amenable to the most appropriate and computationally intensive analyses using site-heterogeneous models of sequence evolution (Lartillot and Philippe 2004). We also employed several strategies to examine the potential for LBA to bias our inferences. LBA has been suspected of influencing phylogenetic placement of several important metazoan lineages including the ctenophores (Philippe et al. 2009; Pick et al. 2010). To monitor the influence of LBA on our analyses, we included several long-branch taxa with non-controversial phylogenetic positions, including the nematodes *Brugia* (Ghedin et al. 2007) and *Caenorhabditis* (C. elegans Sequencing Consortium 1998), the spider mite *Tetranychus* (Grbic et al. 2011) and the larvacean tunicate *Oikopleura* (Denoeud et al. 2010). We also tested the potential of outgroups and locus selection to induce topological artifacts but find no evidence for LBA in any of our analyses. Finally, we examined the possibility that specific categories of genes could differentially support conflicting phylogenetic hypotheses but find no evidence for gene ontology term enrichment between data partitions that support conflicting topologies.

In summary, we recover a phylogeny that is broadly consistent with the recent view of metazoan phylogeny (Dunn et al. 2014). All of the concatenated analyses and locus-selection experiments reported here support the hypothesis of the Ctenophora as sister to the other metazoan species represented by whole genome data, although support for this node varies among analyses. We find no support for the traditional Coelenterata hypothesis, uniting cnidarians and ctenophores as sister clades. A much greater uncertainty exists in the position of the myriapod *Strigamia* relative to other arthropod taxa as selection of data partitions and type of analysis do affect the resulting topology. Our study illustrates an optimized workflow for future analyses of hundreds or thousands of taxa represented by whole genome data.

## Results

### 1. Topology, support and conflict in the 1080-locus dataset

We retained 1,080 individual alignments of putative orthologs following orthology prediction, removal of spurious sequences, alignment, and trimming (see Methods). In total, our data partitions are enriched for 142 gene ontology (GO) terms across the molecular function, cellular component, and biological process categories relative to a reference genome (supplementary fig. S1). The alignments for these 1,080 loci were concatenated into the ‘Total1080 matrix’ that consists of 385,669 amino acid positions at 75.85% occupancy (Table 1).

**Table 1.** Characterization of matrices assembled for phylogenetic inference.

| Matrix name | Number of loci | Length (amino acids) | Missing data (%) | Criteria of locus selection |
| --- | --- | --- | --- | --- |
| Total1080 | 1080 | 385,669 | 24.15 | All loci that after trimming and filtering of paralogs |
| TaxaMin30 | 609 | 199,667 | 20.44 | Loci with at least 30 taxa present |
| TaxaMin33 | 162 | 44,749 | 20 | Loci with at least 33 taxa present |
| TaxaMin35 | 88 | 31,989 | 13.89 | Loci with at least 35 taxa present |
| 60Boot | 55 | 37,682 | 25.92 | Average locus tree bootstrap 60 or more |
| MareMatrix | 143 | 35,030 | 20.02 | MARE (Misof et al. 2013) algorithm filtering with alpha at 3.15 |
| Slow108 | 108 | 33,580 | 18.91 | 10% of the most slowly evolving loci |
| LowLB | 171 | 60,397 | 22 | Low LB scores in the outgroups, sponge, and ctenophore |
| Saturation108 | 108 | 36,954 | 19.78 | 10% of the least saturated loci |
| Best108 | 108 | 41,808 | 15.55 | 10% of loci scoring best in taxon occupancy, saturation, rate of evolution, average bootstrap, and LB scores |
| NoOutgr | 143 | 35,022 | 18.49 | As MareMatrix but realigned without *Monosiga* and *Salpingoeca* outgroups |
| NoAmphi | 143 | 35,032 | 19.75 | As MareMatrix but realigned without the sponge *Amphimedon* |
| NoOutgrAmphi | 143 | 35,009 | 18.13 | As MareMatrix but realigned without the sponge and the outgroups |

We inferred the topology from the Total1080 matrix under maximum likelihood (ML; Felsenstein 1981; Stamatakis 2014) using best-fitting empirical models of protein evolution (Dayhoff et al. 1978; Le and Gascuel 2008) for each partition (Figure 1). The topology of this tree reflects the emerging (Dunn et al. 2008; Hejnol et al. 2009; Ryan et al. 2013; Moroz et al. 2014) but still controversial (Philippe et al. 2009; Pick et al. 2010; Philippe et al. 2011; Nosenko et al. 2013) view of the ctenophores (*Mnemiopsis*) as the sister lineage to all other metazoans including sponges (*Amphimedon*). This topology also recovers all major metazoan clades and many widely-recognized relationships, including Bilateria, the protostome and deuterostome split, a monophyletic Ecdysozoa, the sister relationship between tunicates and vertebrates, as well as the monophyly of the latter. In addition, the positions of several long-branch taxa, which include the nematodes *Brugia* and *Caenorhabditis*, the larvacean *Oikopleura* – by far the longest branch in our whole genome metazoan dataset – and the spider mite *Tetranychus*, are each as expected based on previously published studies (Philippe et al. 2005; Zeng and Swalla 2005; Sharma et al. 2014).

**Figure 1.**
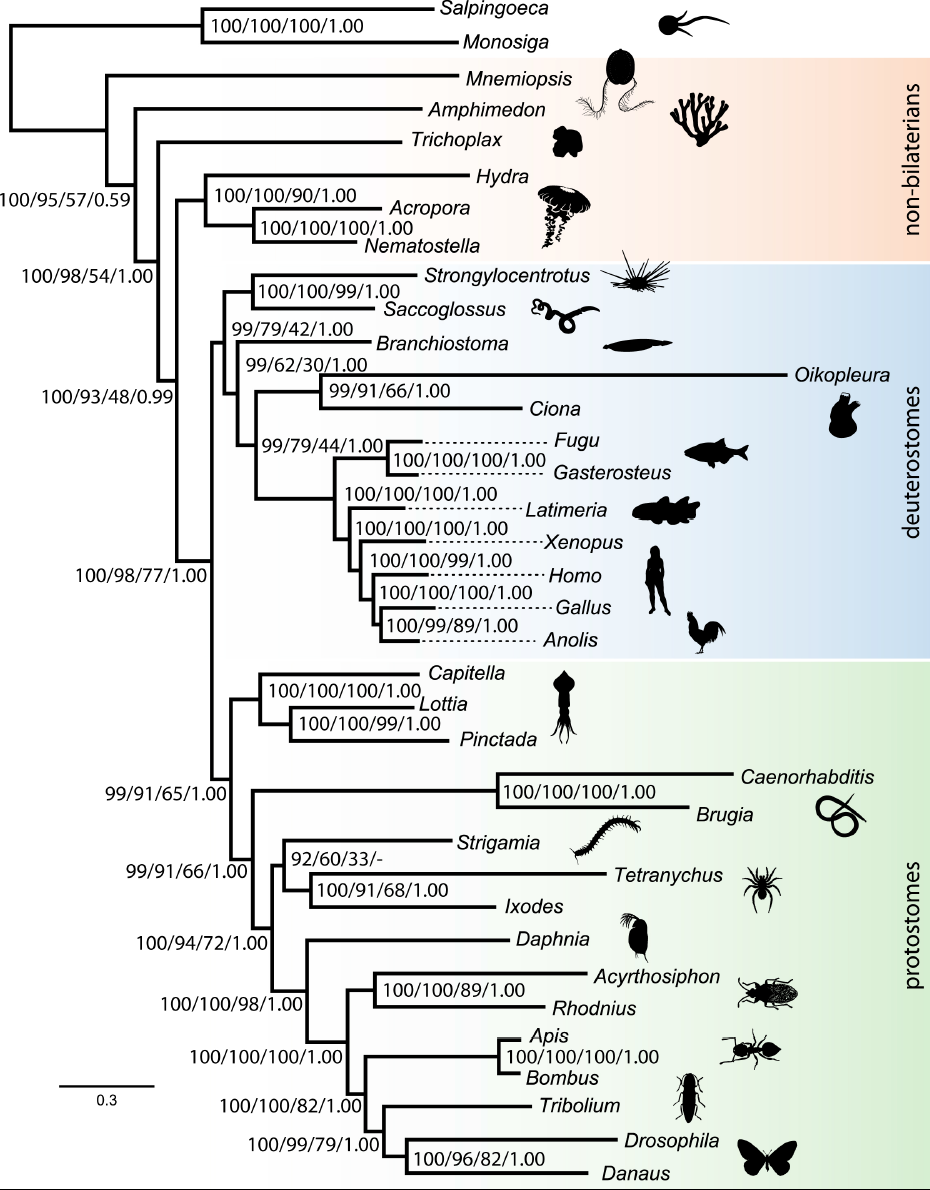
Tree topology and branch lengths from a partitioned analysis of 1080-locus set. Values represent: bootstrap support in Total1080 matrix / 108-locus jackknife from Total1080 locus set / 20-locus jackknife from Total1080 locus set / posterior probabilities from PhyloBayes analysis of Best108 matrix. Scale bar in substitutions per site. Silhouettes from http://phylopic.org. For image attributions see Supplementary Materials.

With the exception of the node joining the centipede *Strigamia* with arachnids at 92%, bootstrap support for the partitioned analysis of the Total1080 dataset was consistently at or near maximum. However, the utility of bootstrap support has been criticized when used on very large data sets (Salichos and Rokas 2013). Jackknife analysis is suitable for assessing the consistency of support across our Total1080 dataset. Among 300 replicates of random datasets of 108 partitions (10% of the total matrix) drawn from all 1,080 loci the support for most nodes in our topology is high, with the exception of some uncertainty in deep deuterostome relationships and in the position of the centipede *Strigamia* relative to other arthropods. Support from 900 replicates of 20-partition jackknife datasets (1.85% of the total dataset) is consistently lower across the backbone of the metazoan tree and deeper deuterostome relationships (Figure 1).

### 2. Topology and support in the Best108 dataset

The Total1080 dataset is too large to analyze under more appropriate, but computationally expensive site-heterogeneous Bayesian models (Lartillot and Philippe 2004). Because of this, we first analyzed each partition separately in order to derive data on 1) information content (Salichos and Rokas 2013; Misof et al. 2013), 2) taxon occupancy, 3) saturation (Philippe and Forterre 1999), 4) long-branch score (Struck 2014), and 5) rate of evolution. We then used these measures to select a set of 108 loci, 10% of the total matrix that scored best across these criteria (see Methods). This ’Best108’ matrix is amenable to computationally intensive analyses and consists of 41,808 amino acid positions at 84.45% occupancy. We also separately assessed the influence of each of these criteria on the phylogeny, as they have each been proposed to potentially negatively impact phylogenetic inference (Baurain et al. 2007; Philippe et al. 2011; Struck 2014) (see supplementary fig. S2 and supplementary fig. S3). Heat maps depicting rate of evolution among partitions for of both the Total1080 and Best108 matrices and the taxon occupancy for each matrix are shown in Figure 2. As with the Total1080 matrix, we performed partitioned ML inference under best-fitting empirical models of protein evolution on the Best108 matrix. In addition, we performed Bayesian analyses on the Best108 matrix under site-heterogeneous CAT-GTR model.

**Figure 2.**
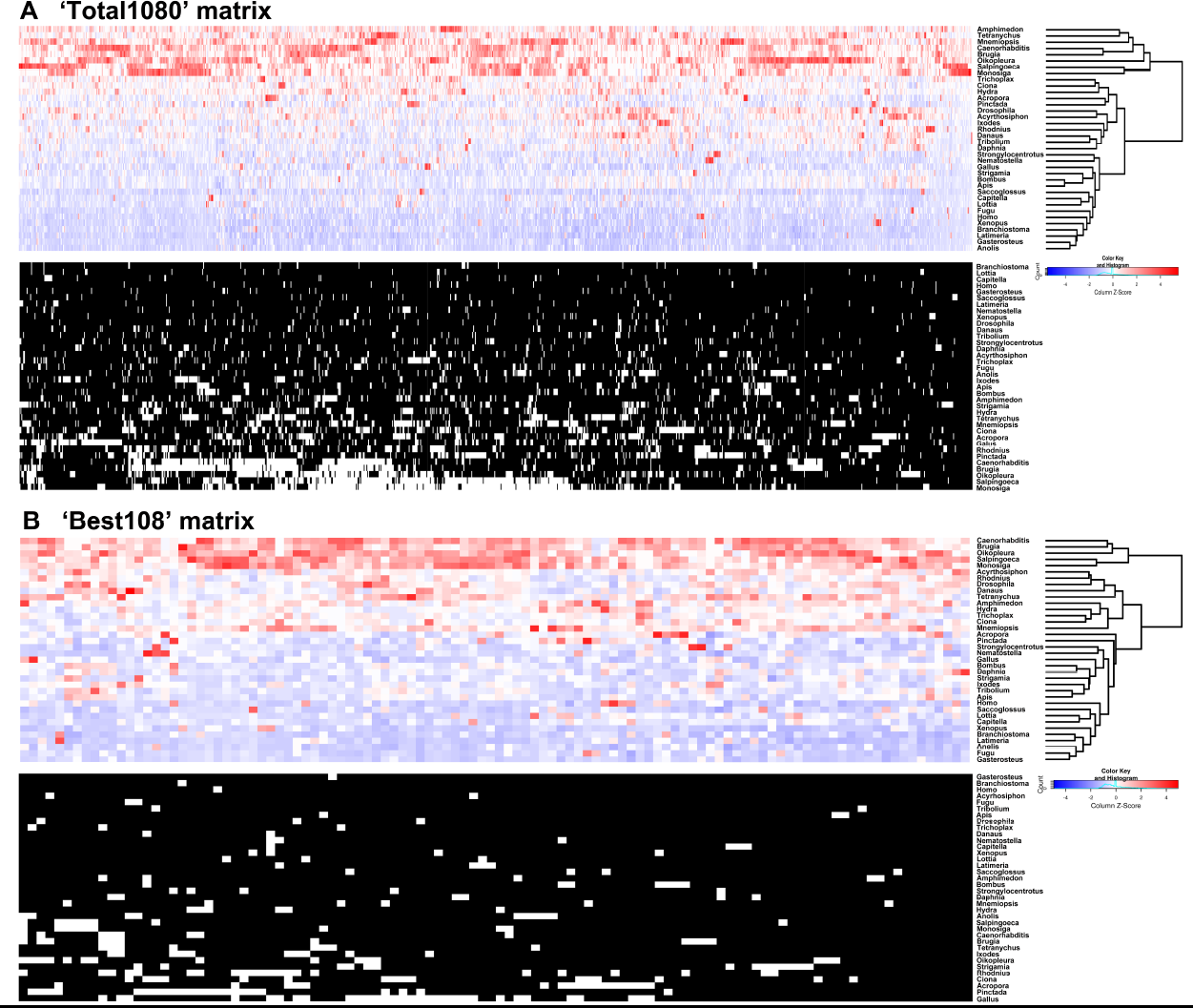
Differences in distribution of long-branch scores (left panels on each side) and gene occupancy (right panels) for ‘Total1080’ and ‘Best108’ matrices. In long-branch score heatmaps, the scores were Z-scaled across columns to highlight among-taxon variability. Red indicates large long-branch scores relative to other taxa and blue denotes low scores. White in gene occupancy plots corresponds to missing data. The cladograms illustrate results of similarity clustering. Note that *Amphimedon* and *Mnemiopsis* do not cluster with long-branched outgroups *Monosiga* and *Salpingoeca* in the ‘Best108’ matrix.

The maximum likelihood (ML) tree for the Best108 matrix shows the same branching pattern as the Total1080 ML tree. The ctenophore is the sister to all other Metazoa in this tree and the centipede is the sister to the chelicerates *Ixodes* and *Tetranychus*. The latter relationship reflects the Paradoxopoda hypothesis (Caravas and Friedrich 2010). However, bootstrap support values for ML analysis of the Best108 matrix are markedly decreased at several nodes in comparison to the 1080-gene matrix. The centipede placement is only weakly supported at 67%, and support for the placement of the lancelet *Branchiostoma* as sister to vertebrates and tunicates is only 80%. However, the ctenophore *Mnemiopsis* is recovered as sister to all other Metazoa in 100% of bootstrap trees in ML analyses of the Best108 dataset (supplementary fig. S2).

Bayesian analyses of the Best108 matrix under the CAT-GTR model produced a topology similar to the ML analysis of the same dataset, with the exception that the position of the centipede *Strigamia* as sister to Pancrustacea now reflects the Mandibulata hypothesis (Snodgrass 1938; Rota-Stabelli et al. 2011) in the Best108 dataset. Interestingly, nodal support given by posterior probability in the Bayesian analysis of this reduced dataset are consistently high across the tree with the exception of the node separating the eumetazoan taxa from the sponge *Amphimedon* and the ctenophore *Mnemiopsis*, which is supported by a posterior probability of only 0.59 (Figure 1).

The CAT-GTR model accounts for the differences in substitution process across sites in a data set, but it does not account for compositional heterogeneity across taxa. This among-branch heterogeneity is present in metazoan alignments from phylogenomic data (Nesnidal et al. 2010) and may also negatively impact phylogeny estimation (Foster 2004; Blanquart and Lartillot 2008). Current implementations of models combining site- and branch-heterogeneity of substitution process are difficult to apply to large data sets (Nesnidal et al. 2010; Boussau et al. 2014). We therefore used an alternative approach that has been shown to be successful in reducing the effects of across-taxon heterogeneity (Nesnidal et al. 2010) and recoded the amino acids in Best108 matrix into six, four, and two categories. We analyzed the recoded Best108 datasets under Bayesian CAT-GTR.

The topology from the data recoded into six categories shows *Amphimedon* and *Mnemiopsis* as sister taxa with a posterior probability of 0.94 and paraphyletic deuterostomes weakly supported at 0.91. This analysis still resolves *Strigamia* as sister to Pancrustacea, but with lower support (supplementary fig. S3D). Recoding the Best108 dataset into four and two categories further confounds resolution, producing trees with non-bilaterian relationships largely unresolved.

### 3. Long-branch attraction

Next we examined the potential for long-branch attraction (LBA) to bias our phylogenetic results. LBA is a phenomenon that causes long-branched taxa to group together artifactually in a phylogeny, often with strong support (Bergsten 2005). LBA is particularly common in datasets with poor taxon sampling or distant outgroups, in which fast evolving ingroup taxa can be ‘pulled’ to the base of the tree by long-branched outgroups. Several studies have indicated that LBA is a potential problem for reconstructing deep animal phylogeny (Lartillot et al. 2007; Philippe et al. 2009; Pick et al. 2010; Philippe et al. 2011; Nosenko et al. 2013). In order to address the potential for LBA to bias our results, we explored various strategies to detect and potentially alleviate the LBA problem.

To test the possibility that the choanoflagellate outgroups affect non-bilaterian relationships through LBA, we assembled three matrices (see Methods) that excluded the outgroups and/or the sponge *Amphimedon*. If the outgroups were to influence the branching order of non-bilaterians, we would expect the internal topology, or the support therein, to be impacted in an analysis excluding the outgroup taxa. Without the outgroups we lose the ability to reliably root the tree, but it is still possible to explore alternative rooting scenarios and to ask if the topology of the ingroup tree is different from those recovered in outgroup rooted analyses. For example, these analyses allow for the examination of possible rooting scenarios where ctenophores are sister to cnidarians, a hypothesis representing the traditional view on non-bilaterian metazoan relationships (Philippe et al. 2009, Nosenko et al. 2013).

We performed partitioned maximum likelihood analysis on the 1) ingroup-only dataset, 2) a dataset where *Amphimedon* was removed and 3) a dataset where both *Amphimedon* and the outgroups were removed (Figure 3). The ingroup-only topology derived from partitioned ML analysis allows for no possible rooting that would place ctenophores and cnidarians together in a monophyletic group (Figure 3A). The sponge-ctenophore bipartition receives 98% bootstrap support, which compares to 100% for all other bipartitions in the tree, except the position of *Strigamia*, which is found in 94% of bootstrap trees and the sister relationship of *Anolis* and *Gallus* at 99% (Figure 3A). If rooted with *Mnemiopsis*, the topology of this tree would be identical to the tree resulting from the ML analysis of a matrix that included outgroups, except that support values are slightly higher in the tree without outgroups (Supplementary Material online, supplementary fig. S2E). If the position of ctenophores were affected by long-branch attraction, we would expect that the removal of outgroup taxa would either alter the branching order or lessen support for non-bilaterian relationships (Philippe et al. 2009). Neither of these possibilities is evident in this analysis. We also examined topologies from partitioned ML analyses in which either the sponge *Amphimedon* was removed (Figure 3C) or both *Amphimedon* and the choanoflagellate outgroups were removed (Figure 3B), and both show a similar pattern. A rooting where *Mnemiopsis* forms a clade with cnidarians, thus supporting the Coelenterata hypothesis, is not possible in any of these analyses.

**Figure 3.**
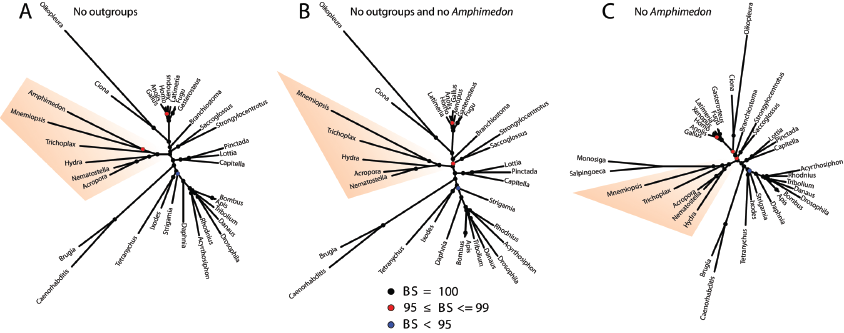
Unrooted trees from analyses with excluding putative long-branch taxa. All analyses were conducted under maximum likelihood, partitioned empirical models. A: Tree inferred without outgroups. B: Tree inferred without outgroups and the sponge *Amphimedon*. C: Tree inferred without the sponge *Amphimedon*. Black circles indicate bootstrap support of 100% from 1000 replicates, red circles indicate support of 95% to 99% and blue circles indicate support of 95% or less. Non-bilaterian metazoan taxa are highlighted.

### 4. Do different genes tell different stories?

Other studies focusing on metazoan phylogeny have suggested that the phylogenetic signal needed to resolve deep relationships is confined to slowly-evolving loci and that specific classes of genes may introduce noise that could mislead analyses (Nosenko et al. 2013; Sharma et al. 2014). In order to explore the influence of rate of evolution of partitions on the support for metazoan relationships, we ranked all loci according to their rate of evolution, approximated by the average branch length of the ML tree inferred for each locus. We then performed a series of unpartitioned ML analyses on matrices that we generated of varying lengths, from few to all loci, beginning with the slowest evolving partitions then progressively adding faster and faster evolving partitions. Unpartitioned ML analysis was conducted for each iteration and support for topologies was assessed using 200 bootstrap replicates. Results from this progressive concatenation approach are detailed in Figure 4.

**Figure 4.**
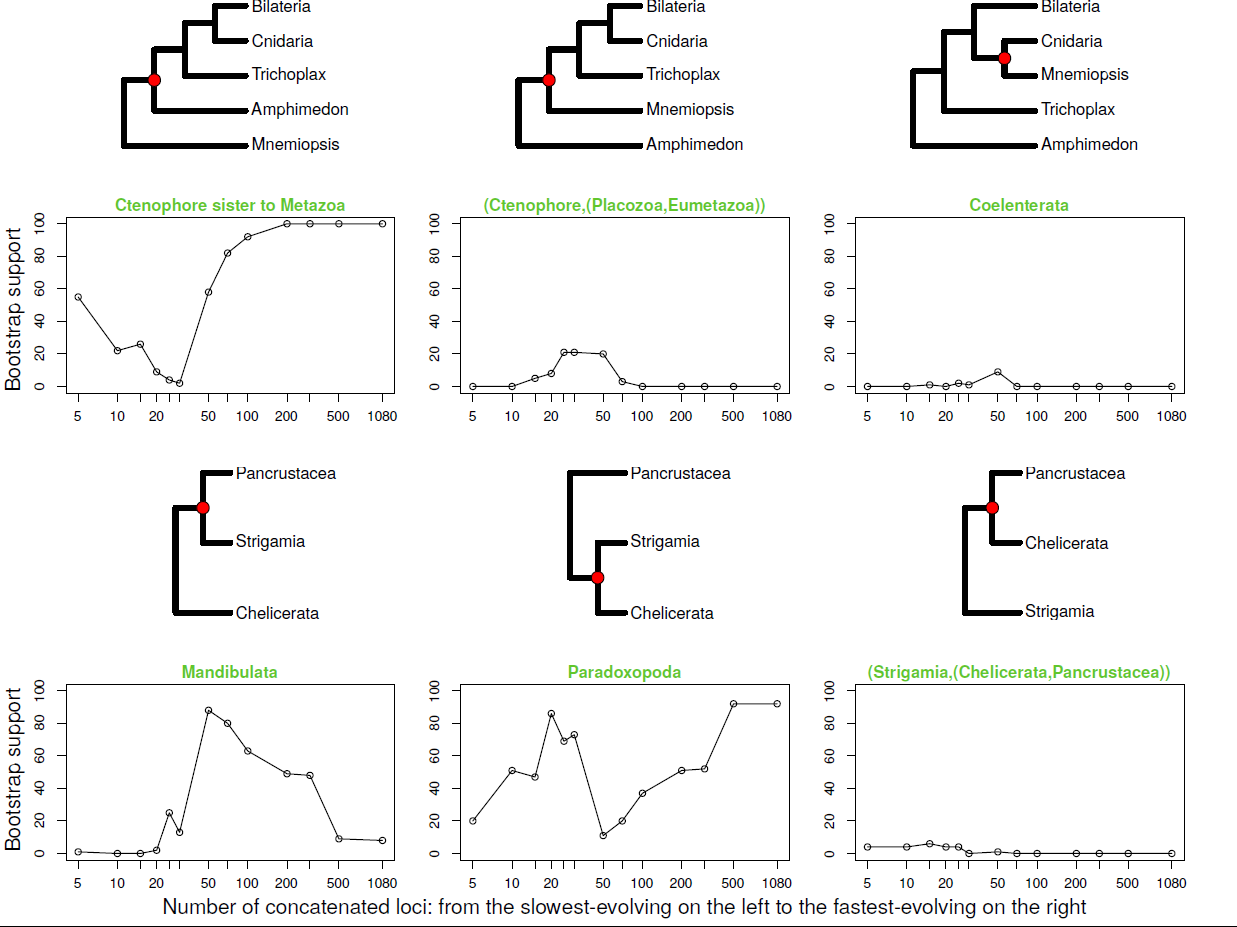
Support for alternative phylogenetic hypotheses under progressive concatenation from the slowest evolving to the fastest evolving loci. The x-axis represents number of loci concatenated in order of rate of evolution, from 5 of the most slowly evolving on the left to including all data at 1080 loci. The y-axis indicates bootstrap support. Red circles in cladograms above corresponding plots denote the node the bootstrap support was measured for.

In each of these analyses, we examined bootstrap support for several possible hypotheses in metazoan phylogeny. These included the following hypotheses for non-bilaterian relationships:

1. Ctenophora sister to Metazoa: ((*Mnemiopsis* (*Amphimedon,* other Metazoa))
2. Ctenophora sister to Eumetazoa: (*Mnemiopsis* (Placozoa, Eumetazoa))
3. The Coelenterata hypothesis, ctenophore as sister to Cnidaria: (*Mnemiopsis* (*Hydra*, *Nematostella*, *Acropora*))

In addition, we compared hypotheses relevant to the placement of the centipede *Strigamia*:

1. The Mandibulata hypothesis, i.e. Myriapod sister to Pancrustacea: (*Strigamia*, pancrustacean taxa),
2. The Paradoxopoda hypothesis, i.e.. Myriapod sister to Chelicerata: (*Strigamia* (*Ixodes*, *Tetranychus*)),
3. Chelicerata plus Pancrustacea monophyletic to the exclusion of the Myriapod: (*Strigamia* (Chelicerata, Pancrustacea)).

Our progressive concatenation analyses show that support for the hypothesis of ctenophore as sister to Metazoa increases rapidly after the addition of greater than 30 partitions. Prior to that number of partitions, support for any of the hypotheses relating to the position of the ctenophores is weak. Following the addition of greater than 30 loci in order of rate of evolution, support for the hypothesis of ctenophore as sister to Metazoa increases to and is maintained at 100% after 200 loci.

The position of *Strigamia* relative to other arthropod taxa is far more complicated in progressive concatenation analyses. Support for the Mandibulata hypotheses peaks at roughly 60 loci and then sharply declines in favor of Paradoxopoda as additional, more quickly evolving loci are added. Support for Paradoxopoda is particularly unusual as it peaks at two inflection points, one after the roughly 20 most slowly evolving partitions are added to the concatenated dataset and the other after 500 partitions are added.

These results suggest the possibility that certain classes of genes, here denoted by their rates of evolution, could differentially favor specific phylogenetic hypotheses. In order to investigate this possibility we conducted a similar analysis where non-overlapping matrices were constructed according to their rank from slowest to fastest evolving. We used a bin size of 108 loci, 10% of the total dataset, per bin (Figure 5, see also Supplementary data).

**Figure 5.**
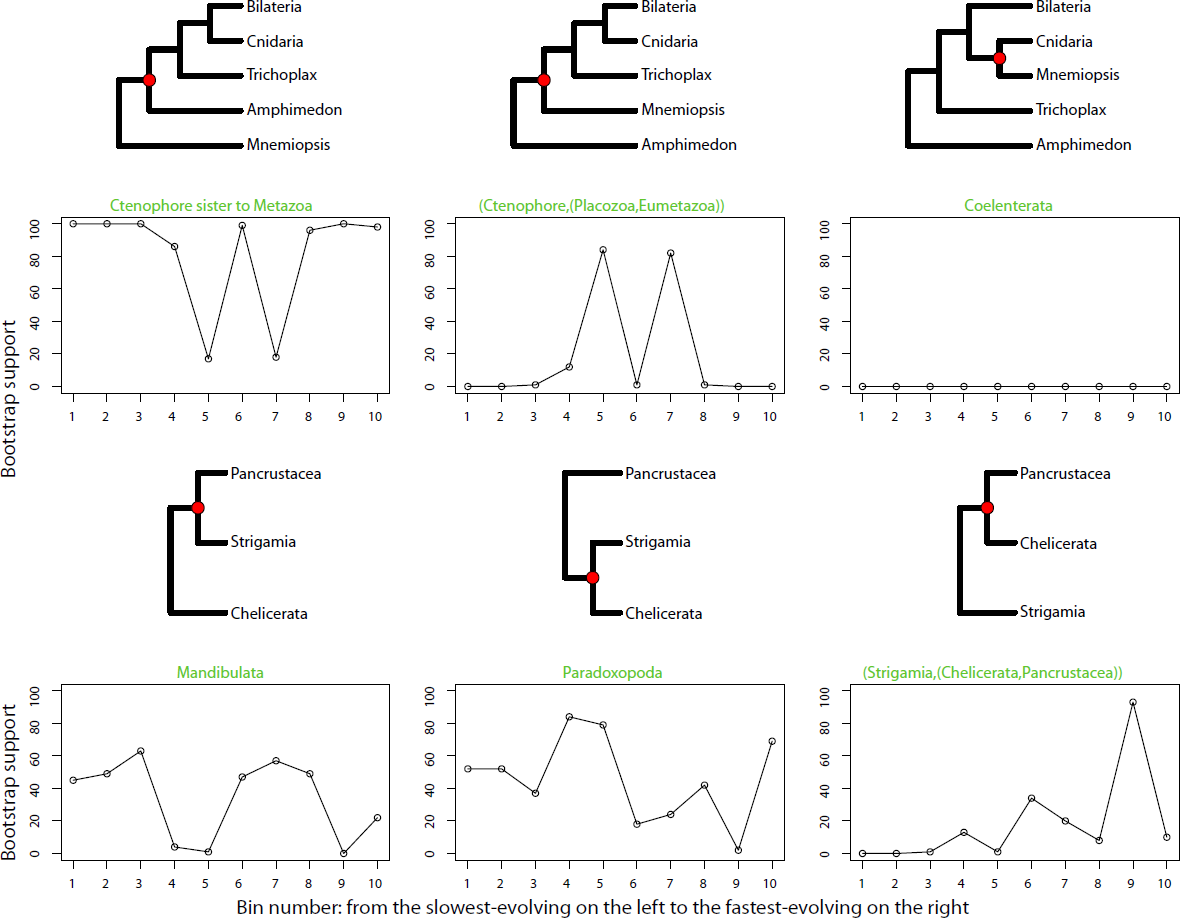
Support for alternative phylogenetic hypotheses across the data. The x-axis represents bin number and the y-axis indicates bootstrap support. Bin number 1 contains 108 slowest evolving loci in the data set and bin number 10 contains 108 fastest evolving loci. Red circles in cladograms above corresponding plots denote the node the bootstrap support was measured for.

In binned analyses, support for the placement of ctenophores as sister to all remaining Metazoa is evident (86-100% of bootstrap trees) in eight out of the ten bins, but was low in bins five and seven (17% and 18%, respectively). We note that the three bins containing the most slowly evolving loci support the hypothesis that the ctenophore is the sister to other Metazoa. The most prevalent competing topology is the one where the sponge is sister to all other metazoans, with ctenophores branching second (Ctenophore (Placozoa, Eumetazoa)). None of the analyses showed consistent support for the Coelenterata hypothesis.

Results from binned analyses were far more complicated regarding the placement of the centipede *Strigamia* relative to other arthropods. Support for either Mandibulata or Paradoxopoda is similar in the first three of the most slowly evolving bins. Support then oscillates between these two hypotheses in the remaining bins. There is also an inconsistent trend toward increased support for the placement of the centipede outside of the Pancrustacea plus Chelicerata clade as the rate of evolution increases.

Our selection of a bin size of 108 loci per bin permitted statistical analyses of GO term enrichment on a bin-by-bin basis. However, these analyses did not reveal a single instance of GO term enrichment in any of the bins compared to the GO terms present in the total dataset. While individual bins may differ in their rates of evolution and the topologies they support, their composition is not significantly different as measured by GO term enrichment.

## Discussion

### Metazoan phylogeny based on genome data

Large data sets are often insufficient to resolve recalcitrant nodes in the animal tree of life and it has long been recognized that simply increasing the amount of data can exacerbate systematic bias in phylogeny estimation (Phillips et al. 2004; Lartillot et al. 2007; Philippe et al. 2011). Because of this, two approaches to improving phylogenomic inference have been proposed. One focuses on the quality of the data and attempts to select only the ‘best’ characters or loci for analysis based on various characteristics of the data (Salichos and Rokas 2013; Misof et al. 2013; Struck 2014). The other is to employ more realistic models of sequence of evolution that account for various systematic biases (Lartillot et al. 2007; Philippe et al. 2011, Roure et al. 2013). Here we leverage both approaches and, due to the large size of our initial data matrix, are able to minimize the impact of various sources of non-phylogenetic signal while retaining a large number of characters for phylogenetic analysis.

One example of how our Best108 dataset represents a refinement of the Total1080 dataset is shown in Figure 2. Here differences in long-branch score and taxon occupancy between the Total1080 and Best108 datasets are compared. In both cases, hierarchical clustering sorts a subset of taxa into a long-branch group of sequences. In the Total1080, this long-branch cluster includes eight taxa including *Mnemiopsis* and *Amphimedon*. In the Best108 matrix, the long-branch cluster is reduced to five taxa and only includes those taxa that reside in non-controversial positions (e.g. both choanoflagellate outgroups, both nematodes and the larvacean *Oikopleura*). In addition, taxon occupancy is enhanced in the Best108 dataset over the Total1080 dataset. For these reasons, we expect that the reduced dataset should contain less phylogenetic noise than the Total1080 dataset.

Results of the majority of our analyses are broadly consistent with the hypothesis for metazoan phylogeny favored in a recent overview by Dunn et al. (2014). The ctenophores emerge as sister to all other multicellular animals and we recover monophyletic Bilateria, protostomes, deuterostomes, and Ecdysozoa. We also find universal support for the recently accepted relationship of the tunicates as sister to vertebrates (Delsuc et al. 2006). Long-branched taxa such as nematodes, the spider mite *Tetranychus,* and the larvacean *Oikopleura* are also universally recovered in uncontroversial positions in our analyses. Most nodes are stable and highly supported across all analyses (Figure 2, see also supplementary figures S2 and S3). This suggests that our whole genome-derived dataset is suitable for the reconstruction of metazoan phylogeny and is robust to the potential influence of long-branch attraction. However, despite high support for most nodes, support near the root of the metazoan tree and for the position of the myriapod *Strigamia* remains variable.

### The root of the animal tree and the position of Ctenophora

Our results are congruent with several recent studies (Dunn et al. 2008; Ryan et al. 2013; Moroz et al. 2014) that depict the ctenophores as the sister lineage to all other metazoans. This hypothesis receives maximum support in all of our analyses conducted under site-homogeneous empirical models of sequence evolution (Figure 1; supplementary figure S2) and our additional analyses suggest that long-branch attraction artifacts do not drive this result (Figure 3). This finding is also consistently supported by progressive concatenation analyses (Figures 4 and 5). However, support for the ctenophores as the sister to the remaining Metazoa is substantially reduced in our analyses inferred under the site-heterogeneous CAT-GTR model (posterior probability = 0.59; Figure 1). Despite this uncertainty, our analyses uniformly support the Parahoxozoa hypothesis, which postulates a single origin of Hox genes in the clade comprised of Bilateria, Cnidaria and Placozoa, to the exclusion of Porifera and Ctenophora (Ryan et al. 2010; Ryan et al. 2013; Moroz et al. 2014). None of our analyses support the Coelenterata hypothesis uniting Cnidaria and Ctenophora, a clade that has been recovered in certain morphological and phylogenomic analyses (Ax 1995; Schierwater et al. 2009; Philippe et al. 2009; Nosenko et al. 2013). In the light of our evidence, we consider the sister relationship of ctenophores to all other animals to be the most conservative interpretation from the analyses presented here, although we recognize, based primarily on weak support from Bayesian analyses under CAT-GTR, that the alternative rooting with sponges as the sister to the remaining Metazoa cannot be rejected.

Among-taxon rate heterogeneity is another potential source of non-phylogenetic signal. We attempted to neutralize this potential confounding factor by recoding our amino acid dataset into fewer amino acid categories and analyzing the resulting datasets under GTR-CAT. Our results were highly inconsistent and failed to resolve several well-accepted clades including a monophyletic Deuterostomia supplementary fig. S3). While these recoding procedures may ameliorate the potential for compositional heterogeneity among taxa, they also drastically reduce the number of informative characters for phylogenetic estimation. This procedure is likely inappropriate for highly conserved datasets such as the Best108 matrix.

### The phylogenetic position of myriapods

The phylogenetic position of myriapods has been considered by morphologists and molecular phylogeneticists alike. The traditional union of the centipedes and milipedes with the clade including crustaceans and arthropods, the Mandibulata, has received support in some recent transcriptome based studies (Regier et al. 2010; Rota-Stabelli et al. 2011; Rehm et al. 2014), but not in others, where myriapods were grouped with chelicerates under Paradoxopoda (clade also dubbed Myriochelata; Dunn et al. 2008; Hejnol et al. 2009; Meusemann et al. 2010).

In our phylogeny, the sole representative myriapod, *Strigamia*, was decidedly the most difficult taxon to place. While Paradoxopoda receives reasonable support in bootstrap replicates of the concatenated Total1080 data set (92%, Figure 2), support for this clade varies drastically across our data and among partitions (Figures 3 and 4). Paradoxopoda is supported in most maximum likelihood trees, but Mandibulata is strongly preferred in a subset of ML analyses and in most of the Bayesian analyses conducted under CAT-GTR (supplementary fig. S2 and supplementary fig. S3). Our findings drawn from whole genome derived data matrices are consistent with previous studies that included much greater taxon sampling and demonstrated the importance of model selection and the potential for LBA artifacts in the placement of the myriapod lineage (Rota-Stabelli et al. 2011). In our analyses, given that the more appropriate model tends to favor Mandibulata, we accept it as the most conservative hypothesis given our data. If correct, this interpretation underscores the importance of systematic bias and data quality in phylogenomic studies (Philippe et al. 2011).

### Conclusions

Our study addresses the problem of basal metazoan relationships using a large dataset drawn exclusively from taxa with publically available whole genome sequences. By applying stringent filtering procedures to our large starting dataset, we were able to obtain a reduced dataset that is still much larger than most previous transcriptome based analyses, but comprised of partitions that that have high taxon occupancy and low potential for non-phylogenetic signal. Results presented here highlight the importance of locus and model selection in phylogenetic analyses. The support for Parahoxozoa is overwhelming in our analyses, suggesting that the traditional grouping of ctenophores and cnidarians in Coelenterata is incorrect. We cannot strongly reject the hypothesis of sponges as the sister lineage to all other metazoans and recognize that more work, particularly in the area of whole genome sequencing, is necessary regarding the root of the animal tree. Much greater uncertainty exists in the position of the Myriapoda relative to other arthropods, but our analyses suggest that this uncertainty can be explained by model selection as previously described (Rota-Stabelli et al. 2011). One obvious drawback of our approach is that taxon sampling is necessarily low compared to other studies that have analyzed transcriptome-based datasets. While numerous workers have emphasized the importance of taxon sampling (Dunn et al. 2008; Philippe et al. 2011), others have emphasized the importance of data matrix size (Rokas and Carrol 2005). Ideally, both parameters would be maximized while maintaining the computational tractability of matrices under the most appropriate models for molecular evolution. Future studies of metazoan phylogeny will benefit from ongoing efforts to sequence the genomes of additional invertebrate taxa that can inform our view of the relationships between the major lineages of animals (GIGA 2014). This is true especially of sponges, where branches subtending this group could be dramatically shortened (Philippe et al. 2009; Moroz et al. 2014) with additional sampling. More genomic resources coupled with better methods that account for systematic biases (Philippe and Roure 2011) and use of additional characters such as presence/absence of genes (Ryan et al. 2013) could soon provide us with a robust phylogeny including all major metazoan lineages (Dunn et al. 2014).

## Materials and Methods

All computational scripts used in the work reported here are found at the GitHub link associated with this study. All sequence datasets, alignments, spreadsheets, annotation files and lists of gene ontology terms for analysis are available at the Dryad link associated with this study. This supplementary data also includes details of all PartitionFinder, RAxML, and PhyloBayes analyses conducted.

*Note to reviewers*: the DOI for the dryad deposition is in progress. A temporary link to these data is here https://www.dropbox.com/sh/6d77yw3li36ib5n/AAD0AKt4WHFQOgr47QzNsR25a?dl=0.

### 1. Taxon sampling and data acquisition

Taxon sampling was aimed to maximize the phylogenetic breadth of species that can inform metazoan relationships, while relying exclusively on species with whole genome sequences. Long-branch attraction (LBA) has been suspected in contributing to the placement of the ctenophores in metazoan phylogeny (Pick et al. 2010). We specifically included other known long-branched taxa such as the nematodes *Brugia* and *Caenorhabditis*, the tunicate *Oikopleura,* and the spider mite *Tetranychus* so that we could monitor the potential for LBA in our dataset. Supplementary table 1 lists these species and the genome databases from which they were obtained.

### 2. Orthology prediction, alignment trimming, and removal of spurious sequences

Gene orthology analysis was performed using a pre-release version 2.0 of the OrthologID pipeline (Chiu et al. 2006). This version of OrthologID uses the MCL algorithm (Enright et al. 2002; Van Dongen 2008) for improved clustering and includes automated extraction of orthologs from gene trees into a partitioned matrix. Amino acid sequences of 1,047,986 gene models from the complete gene sets of all 36 species were used as input to OrthologID, which produced 26,612 orthologous groups with at least 4 species represented. We then selected partitions that included 27 taxa or more for inclusion in our analyses, resulting in a total of 1,162 orthologous groups (OGs). OGs were aligned in MUSCLE (Edgar 2004) using the default settings and trimmed with trimAl v1.4 (Capella-Gutiérrez et al. 2009) using the *-resoverlap 0.5* and *-seqoverlap 50* settings that remove taxa with relatively poor sequence representation within each alignment, followed by *-gappyout* algorithm trimming that removes gap-rich columns. We then conducted maximum likelihood (ML) tree estimation on each locus (see below) and removed potentially spurious sequences with terminal branches more than 5 times longer than the average for the tree. We also discarded partitions that had more than 40% missing data. This resulted in a set of 1,080 curated loci used for further analyses and construction of the ‘Total1080’ data matrix.

### 3. Gene ontology analyses

One randomly chosen gene from each of 1,080 OGs was subjected to blast, annotation and mapping using Blast2GO (Conesa et al. 2005). Gene Ontology identification numbers (GO IDs) for each Metazoan partition were abstracted from this analysis and tested for enrichment against GO IDs from the genome of *Arabidopsis thaliana*, a taxon outside the phylogenetic scope of the focal taxa. Enrichment analyses were performed using Singular Enrichment Analyses and the Fishers Exact Test implemented in agriGO (Du et al. 2010). Enrichment analyses of GO terms between individual bins of metazoan orthologous groups and the total metazoan dataset were also performed using Singular Enrichment Analyses and the Fisher’s Exact Test implemented in agriGO (Du et al. 2010) using GO IDs from the total set of 1,080 OGs as a background annotation.

### 4. Single gene trees, locus selection and construction of the Best108 dataset

We used Phyutility (Smith and Dunn 2008) for concatenation of all multiple-gene matrices and MESQUITE v2.75 (Maddison and Maddison 2008) to convert among file formats.

In order to examine the individual topologies of partitions, we estimated a tree for each of the 1,080 alignments using best-fitting empirical model under maximum likelihood (ML) in RAxML (Stamatakis 2014). We also performed 200 bootstrap replicates for each gene tree. The alignment and corresponding single-gene tree characteristics (see below) served as a basis for several alternative locus selection strategies.

#### a. Locus selection based on information content

We assembled two matrices selecting for information content. One was a concatenation of loci with average nodal bootstrap support higher than 60% (‘60Boot’) (Salichos and Rokas 2013). The other matrix was a result of MARE filtering (Misof et al. 2013) of the Total1080 data set (‘MareMatrix’). We conserved all 36 taxa and used alpha setting of 3.15 (3.00 is default, higher means smaller matrix and higher information content) to obtain a data set of size similar to that of ‘60Boot’ matrix.

#### b. Taxon occupancy and missing data

We concatenated the following matrices with varying level of taxon representation, from 27 minimum taxa per locus to 35 out of total 36: Total1080 matrix that included all 1080 filtered and trimmed orthologous groups with at least 27 taxa represented, ‘TaxaMin30’ with 609 loci having at least 30 taxa, ‘TaxaMin33’ with 162 loci having at least 33 taxa, and ‘TaxaMin35’, a matrix comprising 88 loci with minimum taxon occupancy of 35 out of 36 total species.

#### c. Saturation

We evaluated saturation in each locus by performing simple linear regression on uncorrected p-distances against inferred distances for each locus (Philippe and Forterre 1999). In the absence of sequence saturation, the expectation is that these distances would show a perfect fit to simple linear regression. When there is a need of correction for multiple substitutions, however, the curve will depart from linearity. We used slope and R^2^ of the regression to assess fit in each locus.

#### d. Long-branch score

The so-called ‘long-branch score’ (Struck 2014; LB score) makes it is possible to assess patterns of branch lengths distribution across the data (supplementary fig. S1). This measure is taxon-specific within each locus and defined as the mean pairwise distance of a terminal to all other terminals, relative to average pairwise distance across all taxa. Because of the taxon-specificity, direct comparisons are not possible among loci, and Struck (2014) suggested standard deviation of LB scores as a measure by which loci can be compared. However, we observed that alignments with low standard deviation of LB scores had high proportion of missing data for long-branched taxa. Because of this we implemented an alternative approach, focusing on LB scores of the long-branched *Amphimedon*, *Mnemiopsis*, and the outgroups, *Monosiga* and *Salpingoeca*. We first identified LB mode of density distribution for each taxon, calculated from the Total1080 data set. We then used the number (zero to four) of these focal taxa falling under the mode in each locus to rank all loci. We also concatenated a matrix with 171 loci (‘LowLB’ matrix) with low LB scores and at least three of the four species, therefore minimizing missing data for these target taxa.

#### e. Rate of molecular evolution

We used average branch length of a tree as an approximation of the rate of evolution. A list of loci ranked by average branch length served as a basis for progressive concatenation and binned analyses. We also concatenated a matrix with 10% of the most slowly evolving genes for a partitioned maximum likelihood analysis (‘Slow108’ matrix).

#### f. Construction of the Best108 matrix

We scored the loci by rank in each of the above characteristics (e.g. information content, taxon occupancy, saturation and long-branch score) to assemble the Best108 matrix. We used R packages *seqinr* (Charif and Lobry 2007) and *ape* (Paradis et al. 2004) to compute these statistics, and our R script can be found in the Dryad repository (link) and on GitHub (link). Results are also summarized for each locus in a supplementary table ‘gene_stats.xlsx’ available on Dryad.

### 5. Maximum likelihood analyses of partitioned data sets

We used PartitionFinderProtein (Lanfear et al. 2014) to find optimal partitioning schemes and models for all concatenated matrices. Because PartitionFinder by default uses Neighbor Joining to estimate guide trees, we first inferred a maximum likelihood trees for each unpartitioned matrix using RAxML and used these as user-supplied guide trees for PartitionFinder. We then used RAxML standard versions 8.1 and newer to infer a maximum likelihood tree with support drawn from 1,000 rapid bootstrap replicates.

### 6. Jackknife support in the ‘Total1080’ data set

The jackknife analyses were carried out for 300 replicates of 108-locus (10%) and 900 replicates of 20-locus matrices randomly selected from the 1,080 locus set with a custom Python script. Maximum likelihood tree was estimated for each unpartitioned matrix under the best empirical model selection scheme in RAxML. The support values were then drawn on the concatenated tree using RAxML.

### 7. Bayesian analyses of concatenated datasets

For Bayesian inference, we used PhyloBayes MPI v1.5a (Lartillot et al. 2013), with CAT-GTR as the amino acid replacement model. Analysis with recoded amino acids was performed using PhyloBayes 3.3f (Lartillot et al. 2009). We used three different recoding schemes, which recoded amino acids with six, four, and two groups corresponding to the “dayhoff6”, “dayhoff4”, and “hp” schemes for the *-recode* option in PhyloBayes. Two independent Monte Carlo Markov chains were produced for every matrix. The resulting tree for each matrix is the majority-rule consensus of all trees pooled across both chains sampled at equilibrium. Trace plots were generated using the *mcmcplots* package (Curtis 2012) in R.

### 8. Progressive concatenation and binned analyses

To assess the effect the partitions with high rates of evolution have on the inference, we also incrementally concatenated loci evolving at increasing rates. We sorted the 1,080 gene partitions by their rates of evolution, and created 10 matrices by concatenating 5, 10, 15, 20, 30, 50, 100, 200, 300, and 500 slowest evolving loci. We ran a 200-bootstrap replicates, unpartitioned RAxML search on all these matrices and the all-inclusive matrix of 1080 loci. We also performed binned analyses where loci were concatenated into 60-gene (18 matrices total), 108-gene (ten total), and 216-gene (five total) non-overlapping bins and subjected to a RAxML search as the above. We then mapped bootstrap support for nodes in alternative topologies using RAxML for all progressively concatenated matrices and bins. The trees and support from these experiments can be found in supplementary data on Dryad.

---

**Supplementary figure S1.**

Gene ontology (GO) term enrichment for the Total1080 dataset. GO terms for the genes that comprise the Total1080 dataset were tested for enrichment against the GO term set for the *Arabidopsis* gene model TAIR9 (Zhou et al. 2010). *Arabidopsis* was selected because it is an opisthokont outgroup. When compared to the *Arabidopsis* GO term set, the Total1080 dataset is enriched for 142 GO terms across the Cellular Component, Molecular Function and Biological Process categories.

**Supplementary figure S1.**

Additional trees with branch lengths from partitioned maximum likelihood analyses under empirical models. All support values are based on 1000 bootstrap replicates. A: Tree from 609 concatenated loci that had taxon occupancy of 30 or more; B: Tree from 162 concatenated loci that had taxon occupancy of 33 or more; C: Tree from 88 concatenated loci that had taxon occupancy of 35 or more; D: Tree from 55 concatenated loci with average bootstrap support 60 or higher; E: Tree from 143 concatenated loci selected using MARE algorithm; F: Tree from 108 concatenated loci that were the slowest evolving; G: Tree from 171 concatenated loci where long-branch scores were low in *Amphimedon, Mnemiopsis* and the outgroups; H: Tree from 108 loci that were the least saturated; I: Tree from 108 ‘best’ concatenated loci (Best108 matrix).

**Supplementary figure S2.**

Additional trees with branch lengths from Bayesian analyses under site-heterogeneous CAT-GTR model. All support values are expressed in posterior probabilities. A: Tree from concatenated loci with average bootstrap support 60 or higher; B: Tree from 143 concatenated loci selected using MARE algorithm; C: Tree from 108 ‘best’ concatenated loci (Best108 matrix); D: Tree from Best108 matrix recoded into six amino-acid categories; E: Tree from Best108 matrix recoded into four amino-acid categories; F: Tree from Best108 matrix recoded into two amino-acid categories.

